# The AP2/ERF transcription factor *HcERF5* confers drought tolerance via ABA-mediated signaling in kenaf

**DOI:** 10.1101/2024.08.01.606231

**Authors:** Dengjie Luo, Samavia Mubeen, Muzammal Rehman, Shan Cao, Caijin Wang, Jiao Yue, Jiao Pan, Gang Jin, Ru Li, Tao Chen, Peng Chen

## Abstract

The APETALA2/ethylene response factor (AP2/ERFs) are pivotal in regulating abiotic stress responses in plants. However, the specific role of ERFs in kenaf’s response to drought stress remains unclear. In this study, a transcription factors *HcERF5* was isolated from kenaf and its role in drought stress tolerance was analyzed. HcERF5 was found to localize in both nucleus and cytoplasm and could be significantly induced by polyethylene glycol-6000 (PEG-6000) and abscisic acid (ABA) in kenaf seedlings. In transgenic *Arabidopsis* expressing the *HcERF5* promoter-driven β-glucuronidase (GUS), strong GUS activity was observed in roots, stems, and leaves. Overexpression of *HcERF5* in *Arabidopsis* enhanced seed germination rates under drought or ABA stress and improved drought tolerance in seedlings by increasing antioxidant enzyme activities, whereas *aterf5* knockout lines exhibited the opposite trend. Additionally, *HcERF5* overexpressing *Arabidopsis* showed significantly increased drought tolerance and reduced sensitivity to ABA. Furthermore, virus-induced gene silencing (VIGS) of *HcERF5* in kenaf reduced drought tolerance, as evidenced by decreased antioxidant enzyme activity, increased stomatal aperture, and elevated levels of malondialdehyde (MDA), reactive oxygen species (ROS), and proline under drought stress. RNA-seq analysis further revealed that *HcERF5* directly regulated ABA signaling pathway. Yeast-two-hybrid (Y2H) assays revealed 29 proteins that interact with HcERF5. Among them, the expression of downstream drought stress-related genes *HcPRK*, *HcRD22*, *HcMAP2*, *HcCAB*, *HcCS*, and *HcCCoAOMT3* were significantly reduced in *HcERF5*-silenced plants. Overall, this study highlights the significant potential of *HcERF*5 in enhancing drought tolerance in kenaf.

**Highlight:**

1. *HcERF5* overexpression enhanced drought tolerance in kenaf, however silencing increased drought sensitivity.
2. *HcERF5* regulates ABA synthesis and increases kenaf stomatal conductance and density under drought stress.
3. *HcERF5* regulates plant hormone signal transduction, MAPK signalling, and phenylpropanoid biosynthesis in kenaf under drought stress.
4. Protein interaction revealed HcERF5 interacts with six stress-response genes.

## INTRODUCTION

Drought stress is one of the significant abiotic factors that negatively impacts plant growth and productivity, posing a threat to sustainable crop production globally (Farooq et al., 2017; Wang et al., 2023). Its detrimental effects on crop yield and development are exacerbated by factors such as population increase, global water scarcity, and climate change. Understanding the mechanisms behind drought resistance in crops is crucial for addressing and mitigating the challenges posed by drought stress (Yang et al., 2017). Plants have developed intricate and sophisticated signalling networks to withstand prolonged drought stress (Meena et al., 2017; Lu et al., 2023). These networks encompass a variety of biological processes, including physiological, biochemical, and molecular adaptations. Key responses include enhanced antioxidant production, stomatal closure, osmotic adjustment, as well as hormonal and transcriptional regulation. These responses collectively involve the alteration of thousands of gene expression patterns to protect the normal cellular functions (Udawat et al., 2016; Haider et al., 2017; Udawat et al., 2017; Gong et al., 2020; Challabathula et al., 2022).

Transcription factors families are crucial for signal transduction and the regulation of gene expression, with the ethylene response factor (ERF) genes being a notable subfamily. The ERF family has a unique AP2 domain, making AP2/ERF an important class of transcription factors found in all plants. Based on their structural characteristics, AP2/ERF is categorized into four subgroups: ERF, DREB, AP2, and RAV. These subgroups play vital roles in plant growth and development, morphogenesis, responses to external damage, pathogen resistance, stress responses, metabolic biosynthesis regulation, and plant hormone-mediated regulation (Tang et al., 2005; Wu et al., 2007; Xu et al., 2007; Zhang et al., 2008; Liu et al., 2014; Do et al., 2020; Feng et al., 2020). Plant hormones are important for plant adaptation to adverse biotic and abiotic stress conditions (Khan et al., 2023). Among different plant hormones, abscisic acid (ABA) is a key phytohormone produced in response to osmotic stress, regulates plant growth and development, and is vital for adaptation to various environmental stresses (Raghavendra et al., 2010; Vishwakarma et al., 2017; Yoshida et al., 2019). The regulation of ABA signaling expression can improve plant tolerance (Lim et al., 2015) and ABA-mediated stomatal movement is essential for transpiration regulation under water stress (Kollist et al., 2014; Qiu et al., 2020). Numerous studies have explored the regulation of ABA content by ERF transcription factors during drought stress. For instance, *AtERF4* as a negative regulator of ethylene and ABA signaling pathways, while *AtERF1* enhances drought stress tolerance by regulating stress-specific genes and integrating ethylene, jasmonic acid, and ABA signalling in *Arabidopsis* (Yang et al., 2011; Cheng et al., 2013). In another study, it was found that guard cells overexpressing *AtERF7* exhibited reduced sensitivity to ABA and increased water loss, whereas *AtERF7* RNA interference lines showed high sensitivity to ABA (Song et al., 2005). *AhDREB1*, a member of the ERF5 family, enhances drought tolerance in *Arabidopsis* through the ABA-dependent signalling pathway. Overexpression of *AhDREB1* in *Arabidopsis* increases ABA sensitivity, modifies ABA signaling pathways, and elevates the expression of downstream drought stress-related genes such as *RD29A*, *P5CS1*, *P5CS2*, and *NCED1*. The transcript levels of ABA signaling pathway-associated genes *AtPYL2*, *AtPP2C5*, *AtSnRK2.2*, *AtSnRK2.4*, *AtAREB3*, and *AtABF4* significantly increased under normal growth conditions and after exogenous ABA application (Zhang et al., 2018). Moreover, *OsERF71* regulates genes associated with ABA response and proline biosynthesis in rice. This regulation results in increased sensitivity to exogenous ABA treatment and proline accumulation, thereby improving drought tolerance of plants (Li et al., 2018). In soybean plants, the expression of *GmERF4* is upregulated by cold, salt, and drought but inhibited by ABA (Zhang et al., 2010). In cotton, *GhERF2*, *GhERF3*, and *GhERF6* respond to ABA stress in upland cotton (Jin et al., 2010). Reactive oxygen species (ROS) are highly reactive molecules that function as signalling molecules and induce various stress responses, such as stomatal closure (Medeiros et al., 2020). ROS generated under drought stress interact with ABA to regulate the plant response and enhance its drought tolerance (Geng et al., 2023).

Kenaf (*Hibiscus cannabinus* L.), a member of the Malvaceae family, is an important bast fiber crop globally. It is mainly cultivated in temperate, subtropical and tropical areas of Asia and Africa (Dubois et al., 2013). Kenaf is currently used commercially in over 20 countries, especially in China, Bangladesh, India, and Thailand (FAO, 2021) accounting for 90% of the global kenaf cultivation area and over 95% of global kenaf production (Ding et al., 2016). It is characterized by drought tolerance, salt tolerance, ease of cultivation, marginal land use, high fiber yields, and a variety of industrial applications (Chen et al., 2018). However, the regulatory mechanisms controlling the drought resistance of kenaf are not yet clear and the role of kenaf ERF genes is also poorly understood.

Our earlier transcriptome and qRT-PCR analysis revealed that *HcERF5* was significantly up-regulated in leaves of kenaf seedlings after PEG-induced drought stress, indicating a possible regulatory role of *HcERF5* in drought stress of kenaf (Luo et al., 2023). In this study, to elucidate the role of *HcERF5* in ABA signaling and drought resistance, *HcERF5* was identified and systematically investigated its role in kenaf drought tolerance, via subcellular localization, protein interactions, heterologous overexpression and VIGS analysis. Our evidence suggests that overexpression of *HcERF5* confers ABA insensitivity and enhances drought resistance in plants by enhancing the antioxidant enzyme system. In addition, *HcERF5* affects ABA synthesis and stomatal conductance in *HcERF5* silenced plants under drought stress. The present study provides important and new insights into the drought resistance mechanism of *HcERF5* in kenaf.

## RESULTS

### Identification and characterization of *HcERF5* in kenaf

The coding sequence (CDS) of *HcERF5* is 981 bp in length, encoding a protein of 327 amino acids with an isoelectric point of 5.08 and a molecular weight of 80.8 kDa. Multiple sequence alignment indicated that HcERF5 shares high similarity with HsERF5, GhERF5, and GrERF5, and notably, it has a conserved AP2/ERF domain composed of 58 amino acids (Fig. 1A). Phyre2 predictions suggested that HcERF5 has a single transmembrane α-helix (Fig. 1B). Phylogenetic analysis revealed that HcERF5 is most closely related to Hibiscus syriacus based on amino acid sequences (Fig. 1C). Subcellular localization experiments demonstrated that HcERF5-GFP fluorescence was detected in both the cytoplasm and nucleus, while GFP alone or the nuclear localization marker control showed fluorescence throughout the cell and nucleus (Fig. 1D). These findings indicate that HcERF5 is localized in both the cytoplasm and nucleus.

**Fig. 1.**
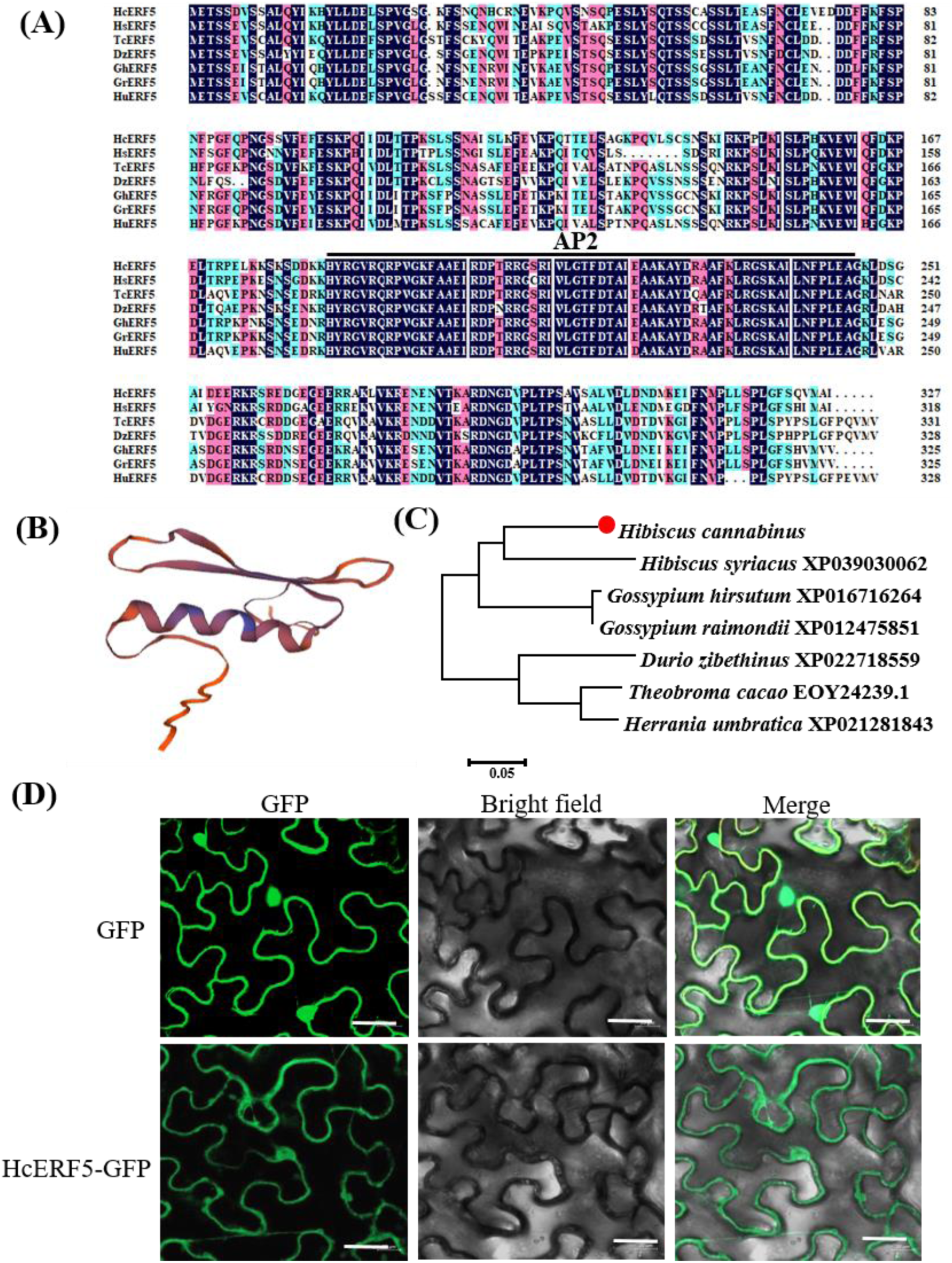
Expression analysis and sequence analysis of HcERF5. (A) Multiple sequence alignment of HcERF5 with its homologous proteins from other plant species. The conservative AP2 domains are overlined in black. (B) Predicted 3D structure of HcERF5 generated using the Phyre2 server. (C) Phylogenetic tree of HcERF5 with its homologous proteins from other plant species. A phylogenetic tree of HcERF5 and its homologous sequences constructed by using the neighbor-joining method using the MEGA 6.0 software. (D) Subcellular localization of HcERF5. Bar: 10μm.

### *HcERF5* can be induced by osmotic stress and ABA treatment

To investigate the role of *HcERF5* in drought and ABA phytohormone signaling, the expression levels of *HcERF5* in kenaf leaves treated with 20% PEG 6000 or 100 μM ABA were evaluated using qRT-PCR. During drought conditions, *HcERF5* expression increased from 2 hours and peaked at 12 hours, reaching 3.1 times the level observed at 0 hours. Although the expression declined after 12 hours, it remained significantly higher than the baseline level before treatment (Fig. 2A). The expression level of *HcERF5* did not show a significant change after 2 hours in ABA treated plants compared to the control group. However, after 4 hours, the expression levels increased significantly, peaking at 12 hours with a 23.2-fold increase from the initial level. With ABA treatment, the expression level of *HcERF5* did not show a significant change after 2 hours compared to the control group. However, after 4 hours, the expression started to increase significantly, peaking at 12 hours with a 23.2-fold increase from the initial level. Following this increase, the expression levels began to decline, but remained 18.2 and 13.4 times higher at 24 and 48 hours respectively, compared to the levels at 0 hours (Fig. 2B). These findings suggest that *HcERF5* is induced and expressed under both drought stress and ABA treatment, indicating its important role in the stress response.

**Fig. 2.**
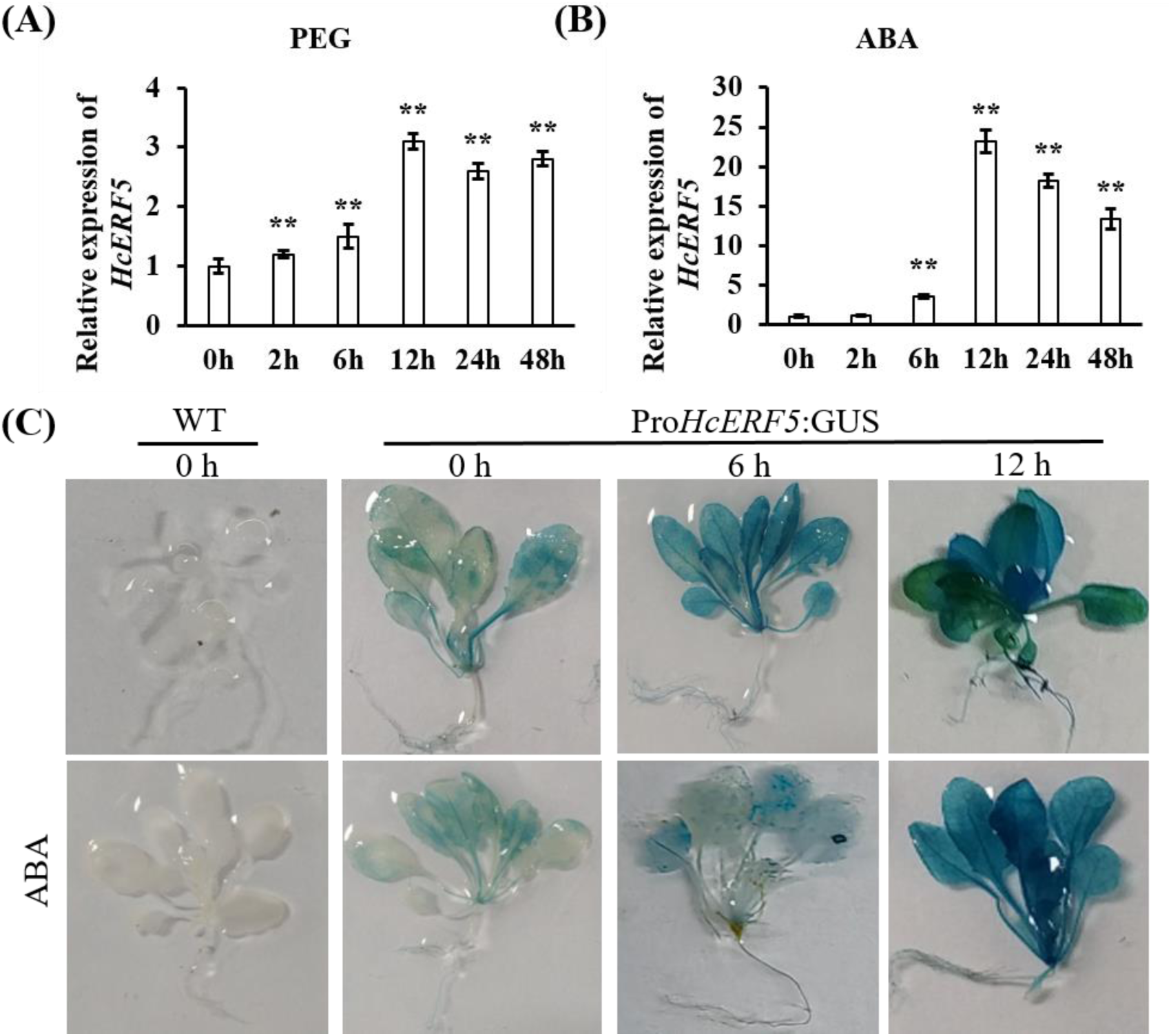
Stress-induced expression assay of *HcERF5*. Expression levels of the *HcERF5* in kenaf leaves under (A) PEG and (B) ABA treatment. The time points of 0, 2, 6, 12, 24 and 48 h were used to observe changes in expression trends with the untreated group at 0 hours serving as the control. Mean and SD were calculated from more than three biological replicates. Asterisks indicate significant differences from control (* for *p* < 0.05 and ** for *p* < 0.01). (C) Analysis of *HcERF5* promoter activity by examining GUS expression in *Arabidopsis* under ABA and drought treatments

The expression patterns of *HcERF5* genes were investigated in the different organ tissues of kenaf. *HcERF5* is expressed in leaves, petioles, stems, and roots, with the highest expression in kenaf leaves, followed by petioles and roots, and the lowest expression in stems (Fig. S2), suggesting that the *HcERF5* have distinct and typical tissue-specific expression patterns. To further investigate the spatial expression of *HcERF5*, the positively transformed Pro*HcERF5*:GUS *Arabidopsis* plants were identified (Fig. S3), and then the GUS activity of 25-day-old plants exposed to 400 mM mannitol or 100 μM ABA for 0, 6, and 12 h was determined. Histochemical staining showed that there is no GUS activity in WT, but in Pro*HcERF5*:GUS *Arabidopsis* plants, GUS activity increases with increasing mannitol treatment time and spreads throughout the plant (Fig. 2C). Besides, GUS activity showed the same change trend under ABA treatment (Fig. 2C). These results indicate that the *HcERF5* gene promoter is highly induced by ABA and mannitol treatments. Collectively, this strongly suggests that *HcERF5* plays a role in ABA signalling and drought stress responses.

### Overexpression of *HcERF5* increased *Arabidopsis* seed germination performance to drought and ABA stress

To elucidate the function of *AtERF5* in *Arabidopsis*, a mutant line named SALK_208574 (referred to as *aterf5*) was identified. In this mutant line, T-DNA was inserted into the fourth exon (Fig. S4A) and genomic DNA PCR was performed to confirm the homozygous lines (Fig. S4B).

*HcERF5* was constructed using the expression vector PBI121 (Fig. S5A) and introduced into wild-type *Arabidopsis* (WT) as background material. The positive seedlings of the T1 generation were screened on the Kan resistance plates (Fig. S5B), and the homozygous T3 generation of the transgenic material was obtained for phenotype analysis. The expression levels of *HcERF5* in the transgenic *Arabidopsis* of the T3 generation and in the *aterf5* mutant were analyzed by semi quantitative PCR. The results showed that *HcERF5* was not expressed in WT and in the *aterf5* mutant of *Arabidopsis*, but only in the overexpressed *HcERF5* lines OE1 and OE2; in contrast, *ATERF5* gene was expressed only in WT (Fig. S5C). Therefore, *HcERF5* was successfully expressed heterologous in *Arabidopsis* and *AtERF5* was silenced in *aterf5* lines.

To investigate the role of *HcERF5* in seed germination under drought and ABA stress, the germination of WT, *aterf5* mutants, and *HcERF5*-OE lines was evaluated over a 7 days period under these stresses (Fig. 3A). The results showed that under normal growth conditions, the germination rate of WT, *aterf5* mutant and *HcERF5*-OE seeds on 1/2 MS media had no obvious differences (Fig. 3B). However, under drought stress, the germination rate of both WT and *aterf5* mutant seeds on 1/2 MS medium supplemented with 200 mM mannitol was slightly delayed (Fig. 3C). When the concentration of mannitol was increased to 400 mM, the germination rate of WT was higher than that of the *aterf5* mutants, but lower than that of the two overexpressing strains, and the germination rate of the *aterf5* mutants remained low even seven day after germination (Fig. 3D). Seed germination of the *aterf5* mutants was significantly delayed on 1/2MS medium supplemented with 2 μM ABA, and only 38.3% and 70.3% of the seeds of *aterf5* mutants germinated on day 3 and 7 respectively, while about 50.1% and 88.6% of the seeds of *HcERF5*-OE germinated on day 3 and 7 and approximately 40.5% and 78.4 % of the seeds of WT germinated on day 3 and 7 respectively (Fig. 3E). Their germination rates showed a similar trend at concentrations of 4 μM ABA (Fig. 3F). It can be concluded that *aterf5* in *Arabidopsis* can increase the sensitivity of *Arabidopsis* seeds to drought and ABA stress compared with WT, while overexpression of *HcERF5* can decrease the sensitivity of *Arabidopsis* seeds to drought and ABA stress. These results indicate that *HcERF5* has a positive effect on seed germination under drought and ABA stress.

**Fig. 3.**
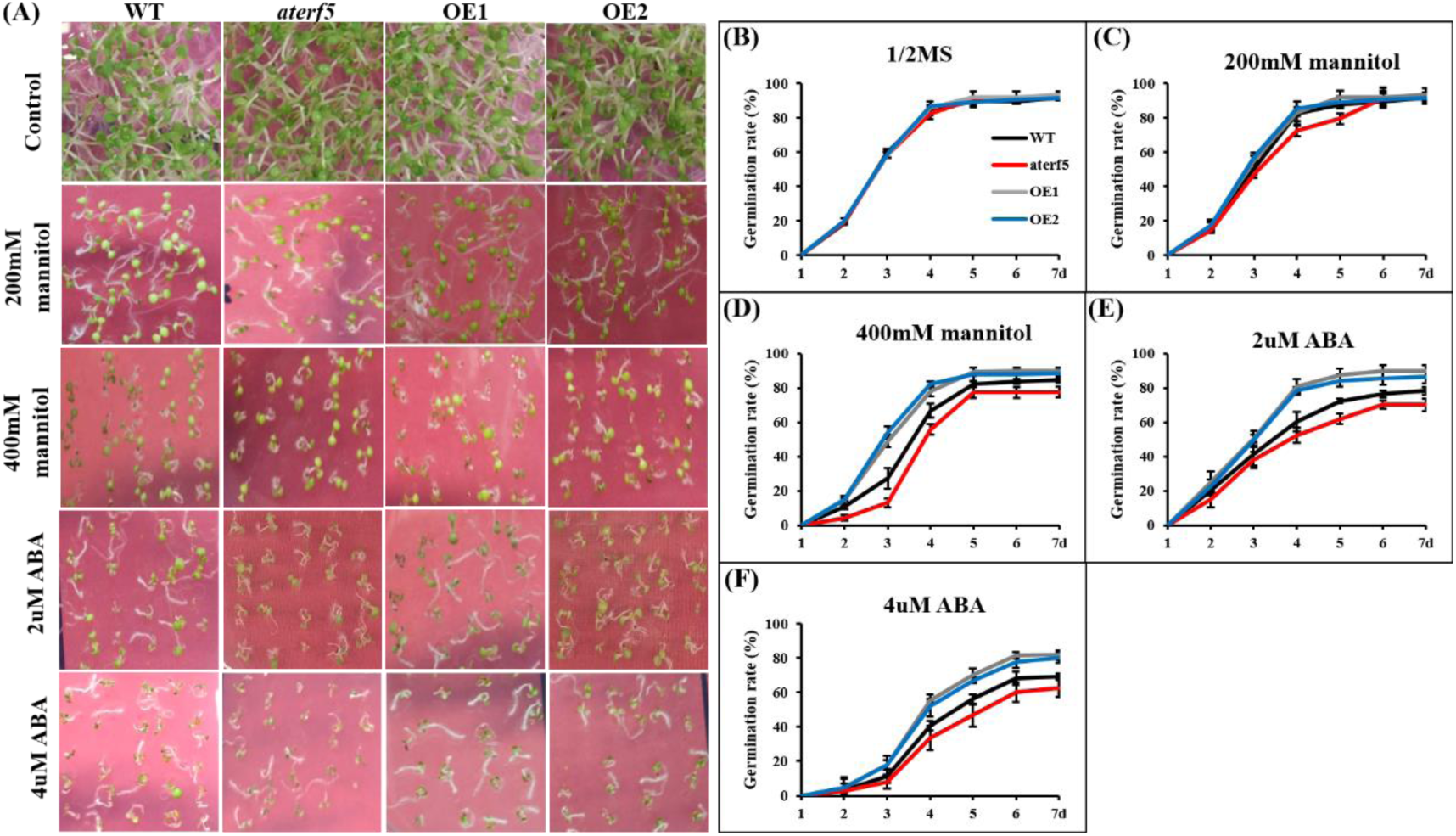
*Arabidopsis* seeds growth on 1/2 MS medium supplemented with different concentrations of mannitol or ABA, and their germination rate. (A) The phenotype of WT, *aterf5* mutant and *HcERF5*-OE lines in different concentrations of mannitol or ABA. (B) Seed germination rate of WT, *aterf5* mutants and *HcERF5*-OE lines on 1/2 MS medium. (C-D) Seed germination rate of WT, *aterf5* mutants and *HcERF5*-OE lines in response to different concentrations of mannitol. (E-F) Seed germination rate of WT, *aterf5 mutants* and *HcERF5*-OE lines in response to different concentrations of ABA. Mean and SD were obtained from three biological replicates.

### *HcERF5* enhances drought tolerance in *Arabidopsis*

To investigate the role of *HcERF5* in root growth under drought stress and ABA treatment, one-week-old seedlings of WT, *aterf5* mutants, and *HcERF5-*OE *Arabidopsis* seedlings were subjected to normal growth conditions, drought stress (200 or 400 mM mannitol in the 1/2 MS medium) and ABA treatment (2 or 4 μM ABA in the 1/2 MS medium). After one week of growth, no significant differences were observed between WT, *aterf5* mutants, and *HcERF5-*OE under normal conditions (Fig. 4A). Under drought conditions, the primary root length of *aterf5* mutants was significantly reduced compared to WT, whereas the *HcERF5*-OE lines exhibited significantly longer root lengths than WT (Fig. 4B). Similarly, the primary root length of the *aterf5* mutants was significantly shorter than WT, while *HcERF5*-OE lines displayed longer roots compared to both WT and *aterf5* mutants (Fig. 4C). These findings suggest that drought stress and ABA treatment markedly influence root length, and that *HcERF5* is a key regulator in the response to these stresses. Additionally, three-week-old Arabidopsis seedlings were subjected to natural drought stress in soil for one week. There was no notable difference between WT and *HcERF5*-OE plants under normal conditions. However, WT and *aterf5* mutant leaves turned yellow and showed severe wilting symptoms under drought stress. Most leaves of *HcERF5*-OE plants remained green and had a higher survival rate after rewatering (Fig. 4D).

**Fig. 4.**
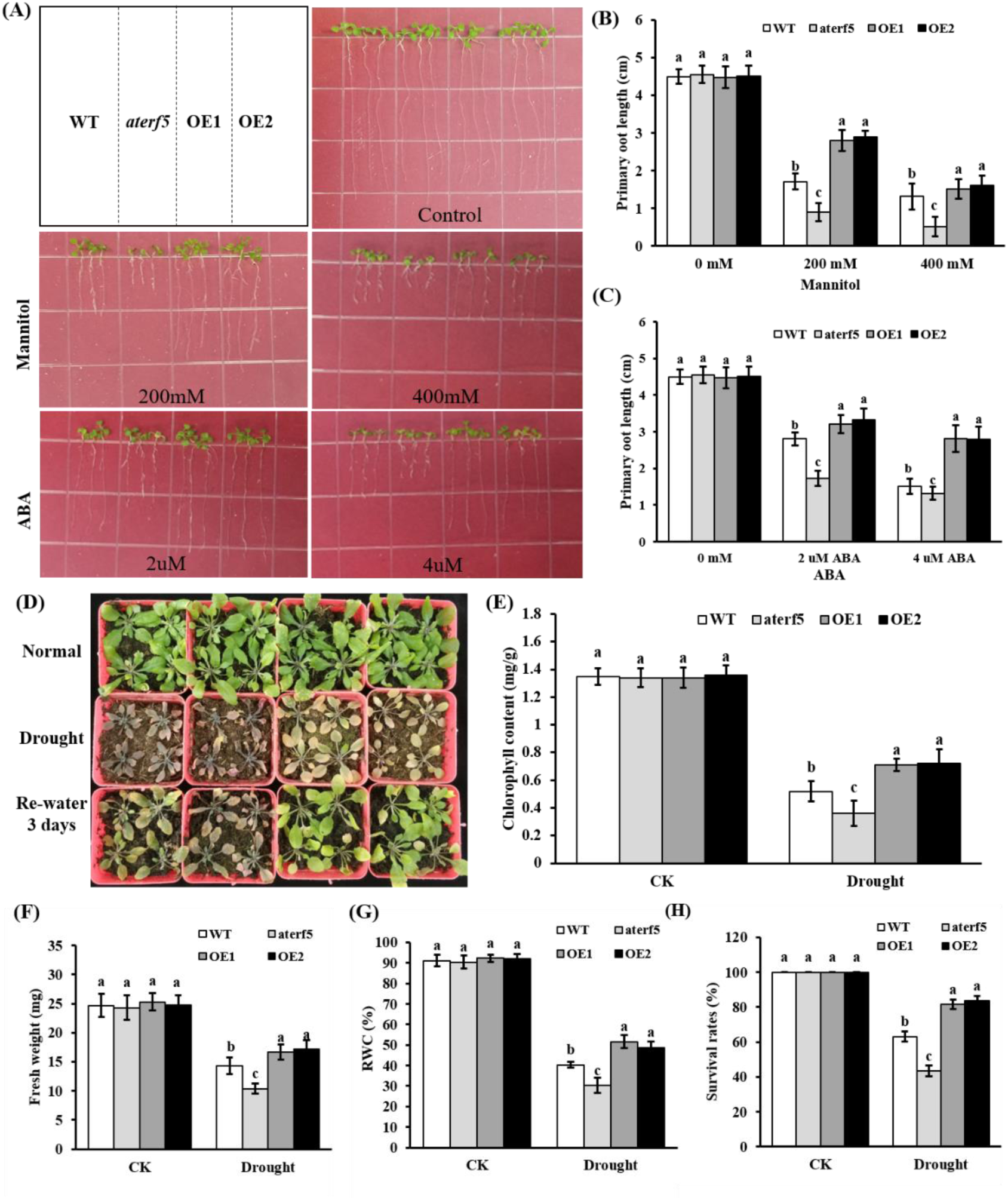
Response of WT, *aterf5* mutants and *HcERF5*-OE *Arabidopsis* plants to drought and ABA treatment. (A) Visualization of root length of WT, *aterf5* mutants, and overexpressed lines under normal, drought and ABA settings. (B-C) Measurement of root length under normal, drought, and ABA conditions. (D) Drought stress and rehydration phenotype. (E) Chlorophyll content. (F) Total fresh weight. (G) Relative water content, and (H) survival rate. Data are shown as the means ± SEs of three biological replicates. Different lowercase letters indicate a significant difference (*P* < 0.05) based on Duncan’s test.

The chlorophyll content in HcERF5-OE under drought conditions was approximately 0.72 mg/g, which was significantly higher than in WT (0.52 mg/g) and *aterf5* mutant (0.36 mg/g) (Fig. 4E). The mean fresh weight of *HcERF5*-OE lines was 17.2 mg, which was significantly greater than that of the WT (14.3 mg) and *aterf5* (10.4 mg) (Fig. 4F). Similarly, *HcERF5*-OE had a considerably higher RWC (51.36%) than the WT (40.4%) or the *aterf5* mutant (30.3%) (Fig. 4G). The survival rate of *HcERF5*-OE lines was 83.7%, surpassing that of WT (63.2%) and *aterf5* mutants (43.5%) (Fig. 4H). The results suggests that *HcERF5-*OE *Arabidopsis* plants displays significantly enhanced drought resistance compared to WT and *aterf5* mutants.

To investigate whether the drought resistance phenotype of the transgenic *Arabidopsis* lines is caused by changes in ROS homeostasis, antioxidant enzyme activity and ROS accumulation were compared in WT, *aterf5* mutant, and *HcERF5-*OE lines under normal and drought stress conditions. The activity of SOD was significantly increased by 96.5%, 67.2%, 124.3%, and 119.1% in WT, aterf5 mutant, OE1, and OE2, respectively compared to the control (Fig. 5A). Similarly, the POD activity was significantly increased by 559.1%, 454.0%, 637.1%, and 608.6% in WT, aterf5 mutant, OE1, and OE2, respectively compared to the control (Fig. 5B). Furthermore, significant increase in CAT activity was 82.8%, 32.2%, 104.5%, and 138.3% in WT, aterf5 mutant, OE1, and OE2, respectively compared to the control (Fig. 5C). Moreover, MDA contents were significantly increased by 70.9%, 95.2%, 26.9%, and 26.3% in WT, aterf5 mutant, OE1, and OE2, respectively compared to the control (Fig. 5D). Quantitative measurements of H_2_O_2_ and O_2_^-^ showed no significant differences in the accumulation of H_2_O_2_ and O_2_^-^ content in leaf tissues of WT and *aterf5* mutants, and *HcERF5*-OE *Arabidopsis* under normal growth conditions. However, after drought treatment, the content of H_2_O_2_ and O_2_^-^ increased significantly in WT and *aterf5* mutants, and *HcERF5*-OE *Arabidopsis*. However, the extent of accumulation in WT and *aterf5* mutants was significantly higher than in *HCERF5*-OE (Fig. 5E and F). Therefore, *HcERF5-*OE *Arabidopsis* plants exhibit lower H_2_O_2_ and O_2_^-^ accumulation than WT and *aterf5* mutant *Arabidopsis* plants.

**Fig. 5.**
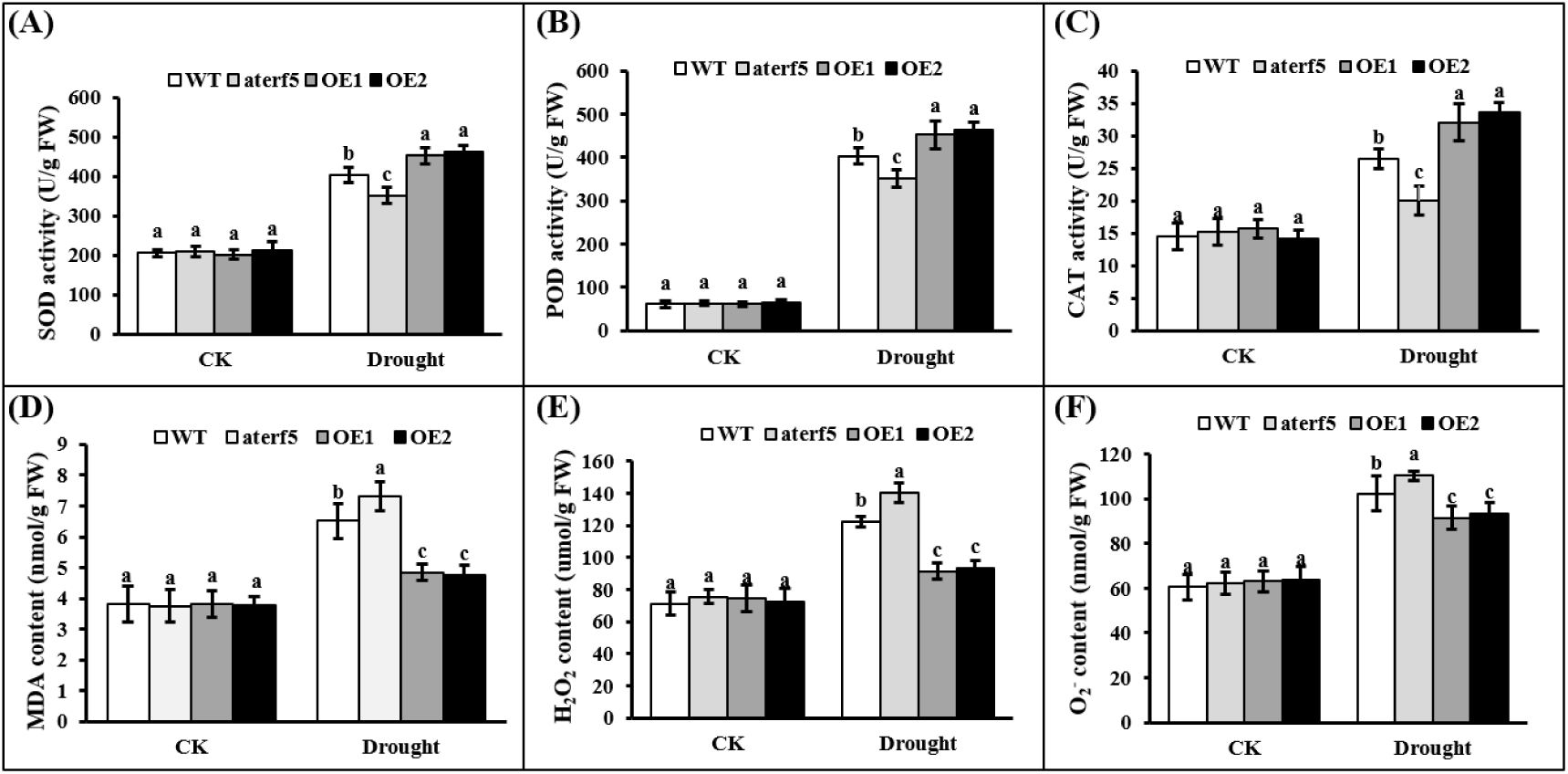
ROS accumulation and activities of antioxidant enzymes under drought stress. (A) SOD activity. (B) POD activity. (C) CAT activity. (D) MDA content. (E) H_2_O_2_ content. (F) O_2_^-^ content. Data are expressed as the means ± SEs of three biological replicates. Different lowercase letters indicate a significant difference (*P* < 0.05) based on Duncan’s test.

### Virus-induced gene silencing of *HcERF5* decreased kenaf drought stress capacity

To further confirm the involvement of *HcERF5* in response to drought stress, *HcERF5* was knocked-down by the VIGS technology. After the successful construction of the recombinant vector pRTV2-*HcERF5* (Fig. S7), the kenaf chloroplast thioredoxin (*HcTrx*) served as positive control, which was used as a reporter gene to detect the status of gene silencing. After 10 days of growth, the *HcTrx* silenced kenaf plants,exhibited a variegated leaf phenotype from the second or third true leaf (Fig. 6A), indicating the reliability of VIGS technology and the down-regulation of *HcERF5* was verified by qRT-PCR in *HcERF5* silenced leaves (Fig. S8). After drought stress, the pTRV2-*HcERF5* silenced seedlings showed severe wilting effect compared with the pTRV2 seedlings (Fig. 6B).

**Fig. 6.**
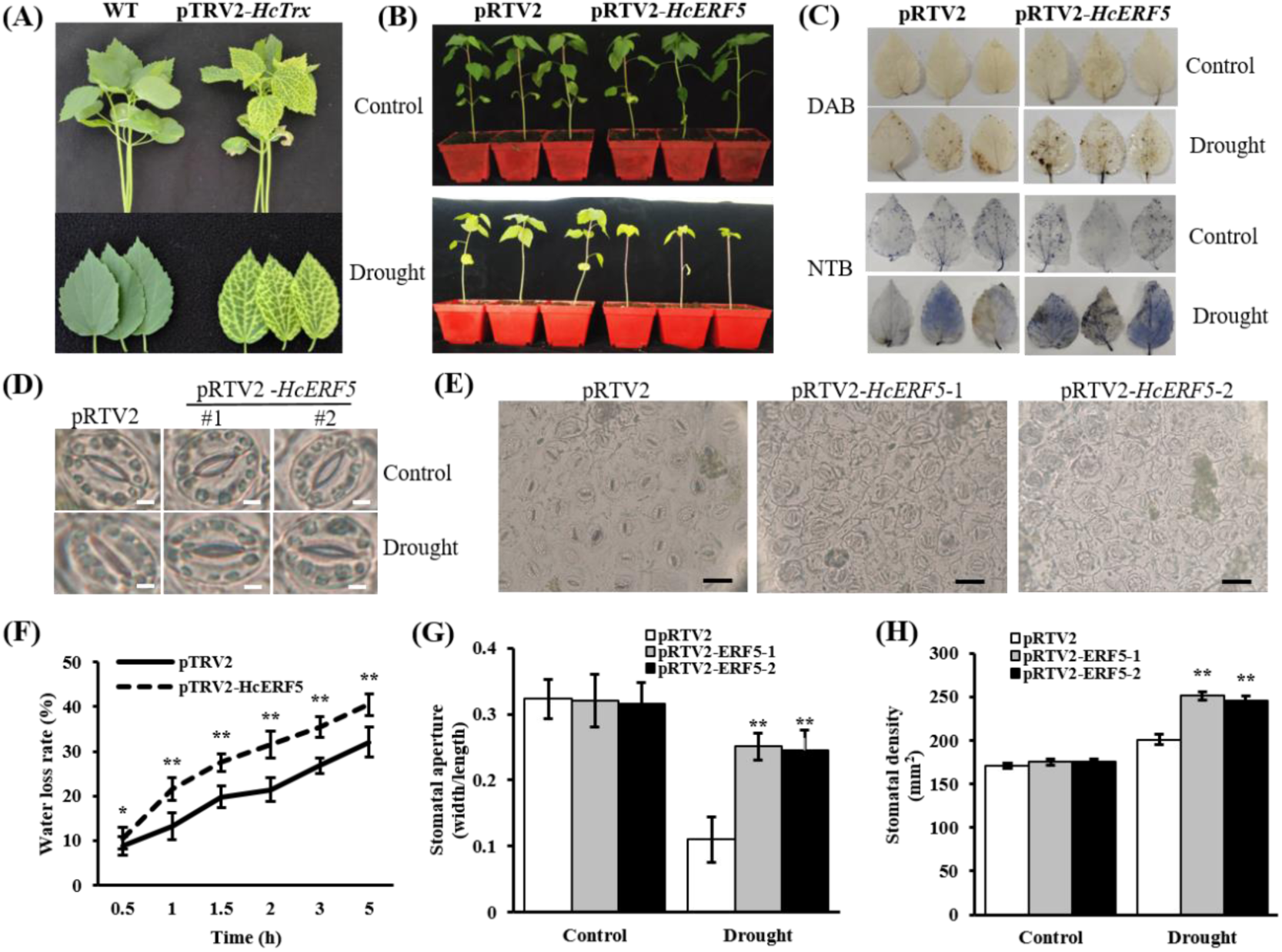
Silencing of *HcERF5* in kenaf reduces tolerance to drought stress. (A) Albino phenotype upon silencing of *HcTrx*. (B) Phenotypes of mock (pRTV2) and VIGS plants (pRTV2-*HcERF5*) under drought stress. (C) H_2_O_2_ and O_2_^-^ accumulation was detected by histochemical staining with DAB and NBT, respectively. (D) Phenotypic analysis of stomata of pRTV2 and pRTV2-*HcERF5* plants. Scale bar = 3 μm. (E) Stomatal density of pRTV2 and pRTV2-*HcERF5* plants photographed under the microscope. Scale bar = 40 μm. (F) Rate of water loss in detached leaves. (G) Measurements of stomatal aperture. (H) Measurements of stomatal density. Data are shown as the means ± SEs of three biological replicates. Different lowercase letters indicate a significant difference (*P* < 0.05) based on Duncan’s test.

DAB and NBT staining revealed that ROS accumulation in pRTV2-*HcERF5* plants was greater than in pRTV2 plants, indicating increased ROS production under drought stress in pRTV2-*HcERF5* plants (Fig. 6C). Furthermore, the rate of water loss in detached leaves and the relative water content of kenaf seedling leaves were examined to determine whether drought-sensitive phenotypes are linked to reduced water retention. The results showed that leaves from pRTV2-*HcERF5* plants exhibited a significantly higher water loss rate compared to control seedlings (Fig. 6F). Since there is a direct correlation between stomatal regulation and leaf water loss, the size and density of stomata on the abaxial epidermis of leaves from both pRTV2 and pRTV2-*HcERF5* plants were analyzed using microscopy under drought conditions (Fig. 6D and E). Measurement of stomatal aperture showed that there was no remarkable difference between pRTV2 and pRTV2-*HcERF5* plants in terms of average stomatal size and stomatal density. However, both the average stomatal size and stomatal density were significantly lower in pRTV2 plants compared with pRTV2-*HcERF5* plants (Fig. 6G and H), indicating that stomatal size and stomatal density are linked to *HcERF5*-mediated leaf water loss. These findings suggested that inhibition of *HcERF5* expression could increase stomatal density and prevent stomatal closure, which promote water loss and reduce tolerance to drought stress.

### Silencing of *HcERF5* increased the accumulation of ROS under drought stress

The plant height and fresh weight of pRTV2-*HcERF5* plants were significantly reduced by 17.2% and 17.9%, respectively, compared to pTRV2 plants after 7 days of drought treatment, although the stem diameter remained unchanged (Fig. 7A and B). The results showed that the activities of SOD, POD, CAT, and GR significantly decreased by 28.8%, 13.1%, 25.7%, and 32.9% respectively, in pRTV2-*HcERF5* plants, compared to pTRV2 control (Fig. 7E-H). However, further quantitative results showed that there was a significant increase of 29.3%, 346.1%, 185.8%, and 47.2% in the content of MDA, H_2_O_2_, O_2_^-^, and proline respectively, in pRTV2-*HcERF5* plants, compared to pTRV2 control (Fig. 7I-L). Overall, silencing of *HcERF5* weakened the antioxidant capacity, and led to excessive ROS accumulation in kenaf, which decreased its tolerance to drought stress, suggesting that *HcERF5* plays a necessary role in kenaf drought-tolerance.

**Fig. 7.**
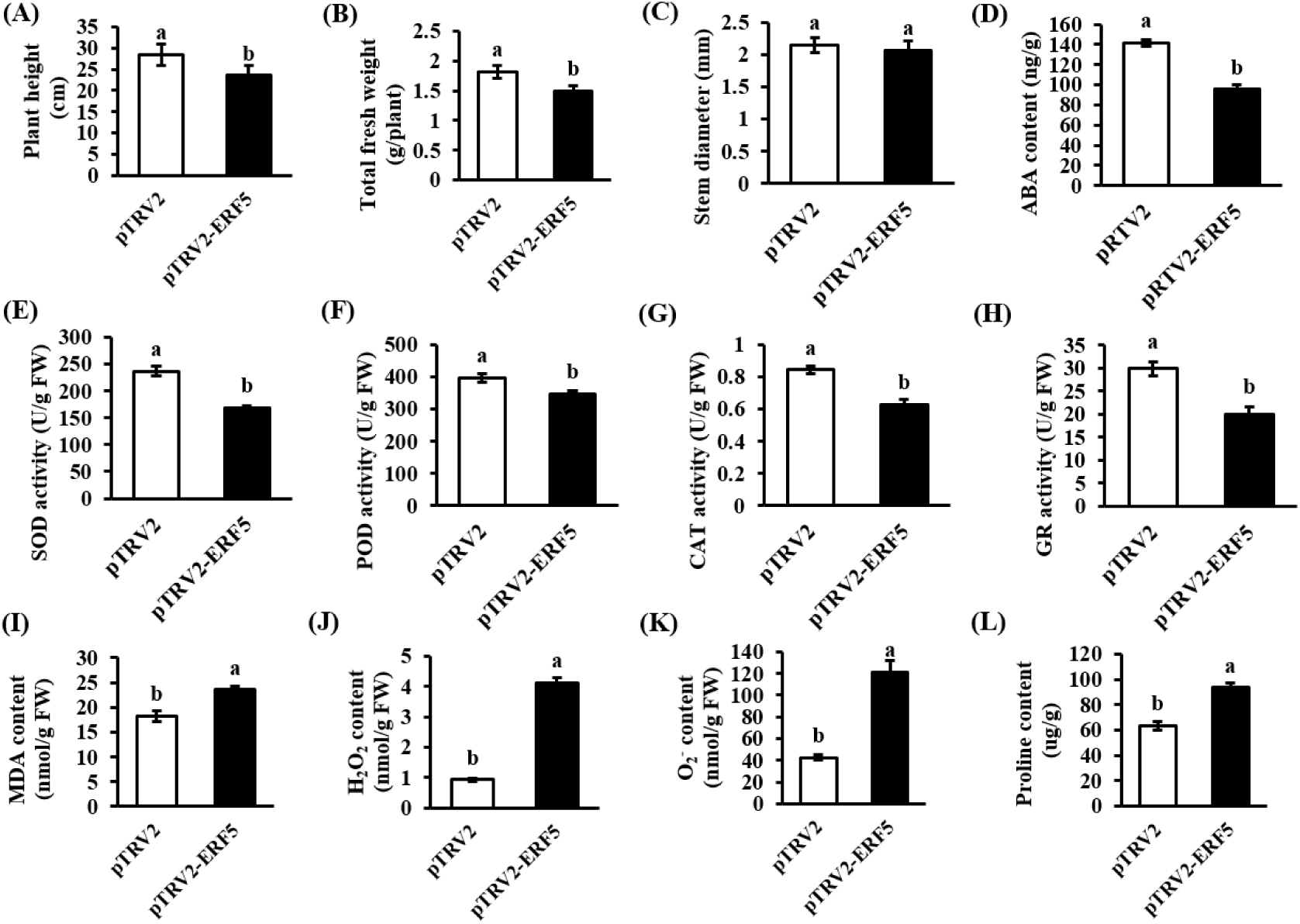
Functional analysis of *HcERF5* under drought stress using VIGS. (A) Plant height. (B) Total fresh weight. (C) ABA content. (E) SOD activity. (F) POD activity. (G) CAT activity. (H) GR activity. (I) MDA content. (J) H_2_O_2_ content. (K) O_2_^-^ content. (L) Proline content. Data are expressed as means ± SEs of three biological replicates. Different lowercase letters indicate a significant difference (*P* < 0.05) based on Duncan’s test.

ABA content and signal transduction are crucial factors in regulating stomatal opening (Jurca et al., 2022). ABA-dependent signal transduction pathways are activated by drought stress (Medeiros et al., 2020). Analysis of ABA content in the leaves revealed a significant decrease in ABA synthesis in pRTV2-*HcERF5* plants under drought stress (Fig. 7D), suggesting that this reduction in ABA content might contribute to the reduced stress resistance observed in *HcERF5*-silenced plants.

### Transcriptome profiling reveals differentially expressed genes regulated by *HcERF5*

To explore the regulatory network of *HcERF5* in response to drought stress, a comparative transcriptome analysis of *HcERF5-*silenced and pTRV2 control plants was performed by RNA sequencing (RNA-seq). The clean reads were submitted with accession number PRJNA1061751 to the Sequence Read Archive (SRA) database. There were 2,489 differentially expressed genes (DEGs) between *HcERF5*-silenced and pTRV2 control plants (|log2FC| ≥ 1 and *p* < 0.05). In the *HcERF5*-silenced line, 722 genes were upregulated and 1767 genes were downregulated in comparison to pTRV2 (Fig. 8A, Table S3). GO enrichment analysis with a *p*-value < 0.05 was conducted on these DEGs to investigate their putative functions. As a result, these DEGs were classified into 45 functional groups, including ‘biological process’ (BP, 19 subcategories), ‘cellular component’ (CC, 12 subcategories) and ‘molecular function’ (MF, 13 subcategories) (Fig. 8B, Table S4). Specifically, 188, 44, 6 and 3 DEGs were annotated with the terms ‘response to stimulus’ (GO: 0050896), ‘antioxidant activity’ (GO: 0016209), ‘detoxification’ (GO: 0098754) and ‘signaling’ (GO: 0023052), respectively. All of which are known to play crucial roles in plant stress tolerance. To gain insight into the biological functions of DEGs in drought tolerance, a KEGG pathway analysis was conducted. This analysis identified 233 DEGs enriched in 14 significant KEGG pathways (*p* value < 0.05) (Fig. 8C, Table S5). These findings suggest that these pathways are crucial for the molecular mechanisms by which *HcERF5* regulates other drought defense genes in kenaf. Notably, 53 genes were enriched in plant hormone signal transduction, 38 genes in the MAPK signaling pathway, and 28 genes in phenylpropanoid biosynthesis. The expression heatmaps of DEGs participating in these three metabolic pathways revealed that these genes were highly differentially expressed (Fig. S9). In addition, the expression levels of genes related to ABA signalling were analysed in *HcERF5-*silenced and pTRV2 plants. Compared to pTRV2, silencing of *HcERF5* altered the expression of several ABA signalling pathway genes, including PYR/PYLs, PP2Cs and SnRK2s (Fig. 8D). In addition, silencing of *HcERF5* also decreased the expression of some genes related to antioxidant enzyme activity, such as POD (Hc.06G002730, Hc.06G003530, Hc.07G015280 and Hc.10G023380), GR (Hc.18G003420), and increased ROS expression (Hc.02G012210 and Hc.03G037930) (Fig. S10).

**Fig. 8.**
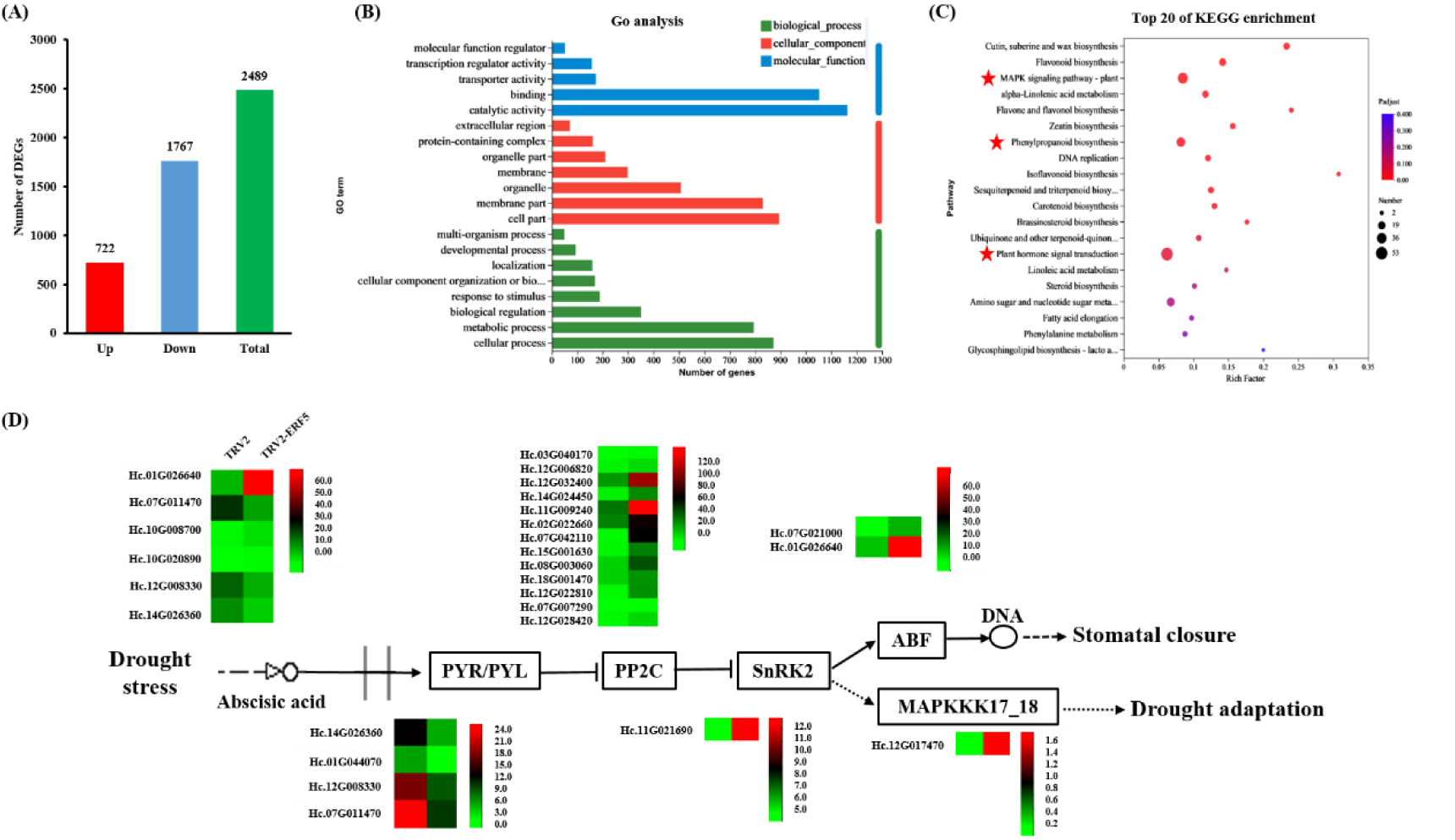
Transcriptome analysis of pTRV2 and pTRV2-*HcERF5* plants under drought treatment. (A) Number of DEGs from *HcERF5-*silenced and pTRV2 plants that are significantly up-regulated and significantly down-regulated. (B) DEGs GO enrichment analysis. (C) DEGs KEGG enrichment analysis. (D) Analysis of gene expression associated with the ABA signaling pathway in pTRV2 and *HcERF5-*silenced plants under drought treatment.

To investigate the different gene expression profiles are regulated by *HcERF5* under drought stress, K-means clustering analysis was performed. The result showed that 2,451 DEGs were distributed among the top 3 K subclusters, accounting for 98.5% of the total DEGs (Fig. S11).

### Screening of HcERF5 interaction proteins

To further investigate the function of the *HcERF5* gene, screening of the HcERF5 gene from a kenaf yeast library was performed. Dot plate analysis revealed that yeast Y2HGold strain containing recombinant plasmids pGBKT7-HcERF5, empty pGBKT7 plasmid, and pGBKT7-53+pGADT7-T was grown normally on SD/-Trp medium, indicating that the expression of recombinant plasmids pGBKT7-HcERF5 in yeast cells is non toxic. At the same time, only the positive control PGBKT7-53+PGADT7-T was detected on SD/-TDO+X-α-gal. The growth on the gal plate and the blue colony indicates that the recombinant plasmid pGBKT7-HcERF5 has no transcriptional activation activity (Fig. 7A). These results indicate that yeast hybridization technology can be used to screen proteins that interact with the HcERF5 protein.

After the mixture of the bait vector pGBKT7-HcERF5 and the library was first screened on SD/-TDO medium, the colonies were further identified on SD/-DDO and SD/-QDO+X-α-Gal. The results showed that the positive control pGBKT7-53+pGADT7-T grown normally on SD/-DDO and SD/-QDO+X-α-Gal medium and turned blue on SD/-QDO+X-α-Gal medium. The negative control pGADT7+pGBKT7-HcERF5 grew normally on SD/-DDO medium, but failed to grow normally on SD/-QDO+X-α-Gal medium without showing blue spots (Fig. 7B). PCR was used to screen the positive strains, and the amplification results showed that most of the bands were about 1000 bp in length (Fig. S12), indicating the effectiveness of screening with kenaf library. These positive clones were sequenced and compared, and unknown proteins were removed. In addition to duplicate clones, 29 proteins that significantly interacted with HcERF5 and had functional annotations were examined (Table S2). According to these annotations, HcERF5 interacted with a number of proteins involved in growth metabolism, stress tolerance, and photosynthesis. These putative proteins have roles in signal transduction or immune processes, indicating that HcERF5 plays an important role in plant stress signal transduction, downstream gene transcriptional regulation, and translation.

To further investigate the interaction between HcCAB and HcERF5. Point-to-point analysis showed that HcCAB interacts with HcERF5 (Fig. 9A). To further verify the interaction between HcERF5 and HcCAB protein, a BIFC method was used based on transient expression in tobacco leaves. The tobacco leaves co-transformed with cYFP-HcERF5 and nYFP-HcCAB showed a clear interaction fluoresence in the plant live imaging system (Fig. S13), and furthermore, a strong yellow fluorescence was detected at the impregnated site, while none was found in the control. This is further demonstrated that HcERF5 and HcCAB can interact in plants (Fig. 9B).

**Fig. 9.**
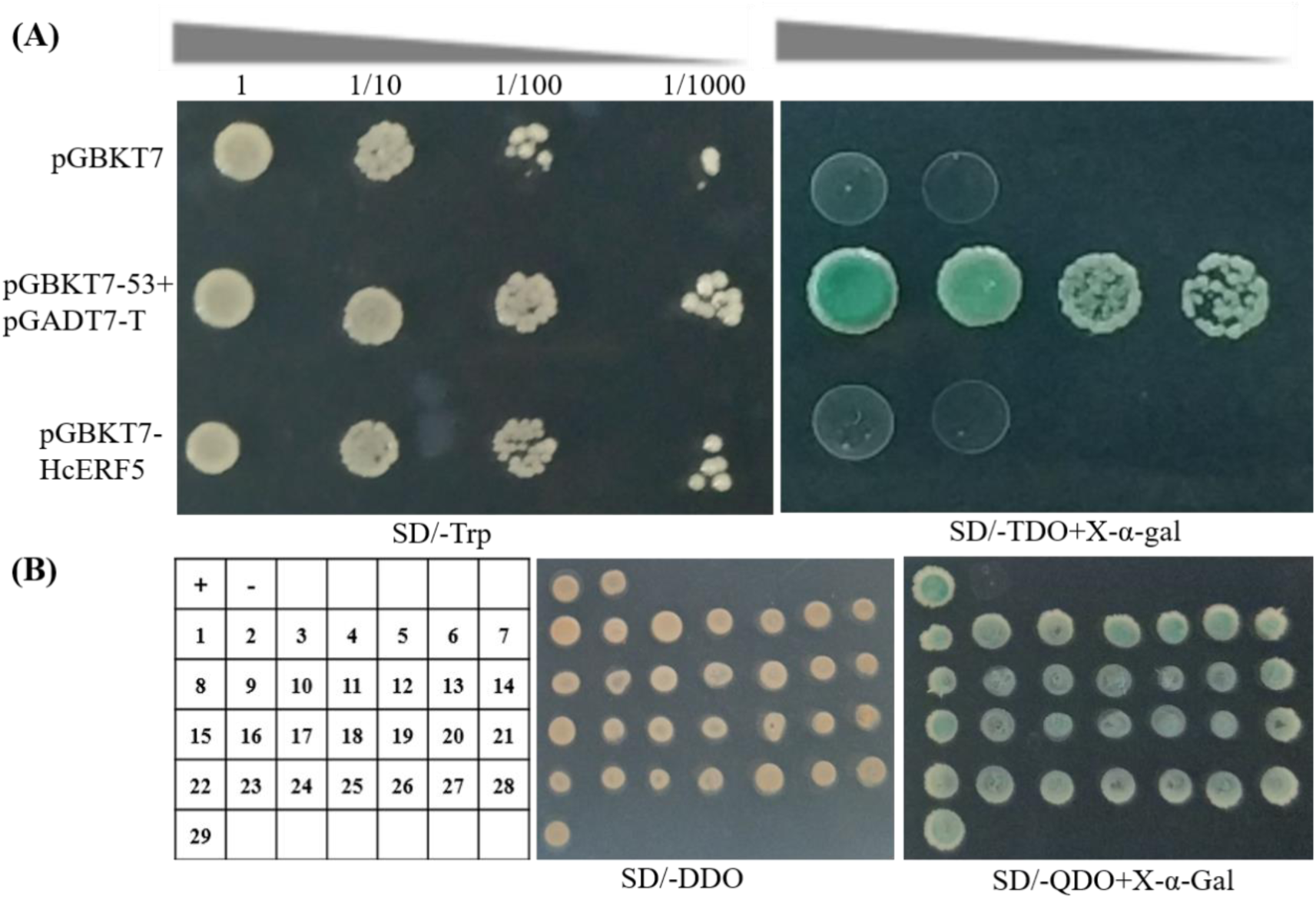
Validation of the interaction proteins for HcERF5. (A) Transactivation activity and toxicity assay of HcERF5 in yeast cells. (B) Validation of interaction proteins for HcERF5. ‘+’ represent pGADT7-T+pGBKT7-53, ‘-’ represent pGADT7-T+pGBKT7-Lam; 1-29 represent interacting colonies with HcERF5. The transformed yeast cells were plated on SD/-DDO, SD/-TDO+ X-α-gal and SD/-QDO+X-α-gal. pGADT7-T+pGBKT7-53 and pGADT7-T+pGBKT7-Lam combinations served as positive and negative controls, respectively.

### Expression of stress-response genes in *HcERF5*-silenced plants

The antioxidant system of the HcERF5-silenced plants was severely damaged, leading to a reduction in plant tolerance to drought stress. To clarify the mechanism, the transcription level of phosphoribulokinase (*HcPRK*), BURP domain protein RD22-like (*HcRD22*), mitogen-activated protein kinase homolog MMK2-like (*HcMAPK2*), chlorophyll a-b binding protein of LHCII type 1-like (*HcCAB*), cysteine synthase-like (*HcCS*) and caffeoyl-CoA O-methyltransferase 3 (*HcCCoAOMT3*) were monitored, based on the interaction genes of the yeast two hybrid. The expression levels of these six genes were significantly lower in the leaves of *HcERF5*-silenced kenaf plants when exposed to salt and drought stress conditions (Fig. 11A-F). These findings imply that *HcERF5* controls the transcriptional activity of these genes to respond drought stress.

## DISCUSSION

### *HcERF5* confer drought tolerance in *Arabidopsis* and kenaf

In recent years, drought stress has significantly impacted crop productivity, making it a serious concern for the sustainability of global agriculture (Morgil et al., 2019). ERFs are key regulators that play a positive role in controlling physiological functions, growth, and stress responses in plants under stress, as part of the ethylene signaling pathway. Various studies have shown that the overexpression of *ERF5* transcription factors can enhance resistance in a variety of plant species (Fujimoto et al., 2000; Fischer and Dröge-Laser, 2004; Jin et al., 2009; Chuang et al., 2010; Moffat et al., 2012; Pan et al., 2012; Son et al., 2012; Dubois et al., 2013; Severo et al., 2015; Li et al., 2023). However, the role of the ERF5 gene from kenaf in drought tolerance has not been well explored. In this study, *HcERF5* gene was cloned, which has a typical conserved AP2/ERF domain and subcellular localization showing its expression in the cytoplasm and nucleus (Fig. 1). Therefore, we hypothesized that it has a similar function to other ERF5 genes. The changes of *ERF5* expression in *tamarix hispida*, tomato and cotton were significantly induced by drought or ABA (Jin et al., 2009; Pan et al., 2012; Liu et al., 2014). In this study, *HcERF5* expression in kenaf was significantly upregulated by drought and ABA, and high expression levels in the leaves, petioles, stems, and roots of transgenic *Arabidopsis*, indicating a potential role in the development of these organs (Fig. 2). The heterologous expression of *HcERF5* in *Arabidopsis* was found to enhance seed germination rates under drought stress and decrease seed sensitivity to ABA. Conversely, *aterf5 Arabidopsis* mutants exhibited reduced seed germination rates under drought stress and increased seed sensitivity to ABA (Fig. 3). Heterologous expression of *HcERF5* in *Arabidopsis* can significantly enhance the drought resistance of seedlings, while the *aterf5* mutant of *Arabidopsis* significantly reduced drought tolerance (Fig. 4). It is well established that increased lipid peroxidation exacerbates cellular oxidative damage in plants under drought stress (Tang et al., 2013). This phenomenon results from the excessive accumulation of ROS such as H_2_O_2_ and O_2_^-^ radicals. Antioxidants are essential for mitigating oxidative damage induced by drought (Wu et al., 2015). Under drought conditions, the levels of H_2_O_2_ and O_2_^-^ in the leaves of the *aterf5 Arabidopsis* mutant were significantly higher than those in WT, whereas WT plants exhibited higher levels of H_2_O_2_ and O_2_^-^ than transgenic plants (Fig. 5E and F). This suggests that the transgenic lines experienced slightly less cellular oxidative damage compared to WT plants. The higher MDA content in plants indicates higher degree of lipid peroxidation and consequently, more extensive cell membrane damage (Sun et al., 2014). Under drought stress, the *aterf5 Arabidopsis* mutant showed the highest MDA levels in its leaves, followed by the WT, with the transgenic plants having the lowest levels (Fig. 5D), indicates that the *aterf5 Arabidopsis* mutant experienced the most severe damage, followed by the WT, with the transgenic plants showing the least damage. Additionally, the activities of SOD, POD, and CAT, which are crucial for ROS degradation, were measured. *Arabidopsis* plants under drought conditions exhibited a significant increase in SOD, POD, and CAT levels. However, the enzyme activity was significantly higher in transgenic plants compared to WT plants, and WT plants had higher enzyme activity than the *aterf5 Arabidopsis* mutant (Fig. 5A-C). This demonstrates that overexpression of *HcERF5* can mitigate cellular oxidative damage under stress by enhancing ROS scavenging capacity. Mutation of *AtERF5* in *Arabidopsis* led to reduced antioxidant enzyme activity, resulting in decreased drought resistance. These findings suggest that *HcERF5* enhances drought tolerance by regulating ROS scavenging.

The growth and development of *HcERF5*-silenced plants were significantly inhibited under drought stress, including plant height and fresh weight (Fig. 7A and B). At the same time, there was a significant decrease in antioxidant enzyme activities (Fig. 7E-H), and increase in MDA, H_2_O_2_, and O_2_^-^ contents (Fig. 7I-K). Consistent with our observation, increased ROS accumulation was recently reported in *HcERF4*-silenced kenaf plants under drought stress (Yue et al., 2022). In addition, accumulation of proline under abiotic stress protects cells from damage by osmotic regulation and free radical scavenging (Kollist et al., 2014). Proline can serve as an indicator of the degree of stress-induced damage (Auriga and Wrobel, 2018), and silencing of *HcERF5* resulted in a significant increase in proline content (Fig. 7H), indicating that silencing of *HcERF5* led to severe drought damage in kenaf plants. Notably GO enrichment analysis of transcriptome DEGs revealed that there is a wide array of FB processes, including immune system processes, detoxification, signaling, and response to stimuli etc. (Fig. 8B, Table S4). The expression of some antioxidant enzymes or genes also changed significantly (Fig. S10). Taken together, this means that *HcERF5* can regulate the expression of related genes, and then improve the physiological indexes such as antioxidant enzyme activity of plants under drought stress, and is a factor that responds positively to drought tolerance.

### *HcERF5* is involved in regulating the ABA signaling pathway affects ABA content and regulates stomatal conductance

ABA plays a pivotal role in plant responses to abiotic stresses, including drought (Hirayama and Shinozaki, 2007; Liu et al., 2023). Under challenging conditions such as drought and high temperatures, there is a notable increase in ABA levels within plants. This elevation triggers the rapid activation of the ABA signalling pathway, which in turn enhances the plant’s ability to withstand stress (Cutler et al., 2010; Wang et al., 2023). A substantial number of transcription factors have been documented to respond to osmotic stress through ABA-dependent manner (Jardak-Jamoussi et al., 2016; Chen et al., 2022). The ABA signal transduction process is the most widely researched and earliest known signaling pathway involved in PP2Cs. In higher plants, the ABA signaling module mainly consist of three parts, namely ABA receptor PYR/PYL/RCARs, PP2C members of A subfamily, and SnRK2s (Fujii et al., 2009). Protein phosphatase 2C (PP2Cs) serve as a key regulators in plant responses to abiotic stresses. In *Arabidopsis thaliana*, PP2CA acts as a central regulator of ABA signaling, negatively influencing plant growth, development, and responses to various stresses (Baek et al., 2019). Our study identified a total of 13 PP2C transcription factors that were significantly up-regulated in pTRV2-*HcERF5* plants (Fig. 8D), highlighting their crucial role in kenaf’s response to drought stress.

Stomata play an important role in plants by regulating water loss and gas exchange. During drought periods, there is a positive correlation between stomatal density and the transpiration rate. ABA influences stomatal conductance by inducing stomatal closure, thereby reducing transpiration water loss and triggering the expression of drought-responsive genes. This mechanism ultimately enhances drought resistance in plants (Lim et al., 2015; Saradadevi et al., 2017). Under drought stress, overexpression of AP2/ERF gene, *IbRAP2-12 Arabidopsis thaliana* lines can upregulate ABA signaling genes, significantly increase ABA content and decrease water loss rate, thereby enhancing the plant tolerance (Li et al., 2019). Our findings are similar to those of previous studies, particularly in observing stomatal closure in both pTRV2 and pTRV2-*HcERF5* plants when subjected to normal and drought stress conditions. Under normal conditions, stomata in both pTRV2 and pTRV2-*HcERF5* plants remained open, with no significant difference observed in the stomatal width-to-length ratio between the two plant types. However, stomatal closure was more pronounced in pTRV2 plants under drought than in pTRV2-*HcERF5* plants (Fig. 6D and G). The changes in stomatal density showed the same trend (Fig. 6E and H). These results demonstrate that *HcERF5* promotes stomatal closure and stomatal density in response to drought, thus reducing the rate of water loss (Fig. 6F). ABA content under drought stress was also measured and found that the accumulation of ABA significantly increased in both pTRV2 and pTRV2-*HcERF5* plants after drought treatment, and the ABA content in pTRV2 plants was higher than that in pTRV2-*HcERF5* plants (Fig. 6G). These findings showed that ABA synthesis under drought is inhibited by silencing of *HcERF5*. Therefore, it is speculated that *HcERF5* enhances drought resistance through ABA signaling.

### *HcERF5* interact with stress-responsive related genes in drought tolerance

Plants have evolved sophisticated protective systems to recognize and respond to adverse environmental conditions, such as the expression of stress-related genes (Nevo et al., 2010). For instance, the expression of PRK in rice is regulated by ABA, gibberellin (GA) and methyl jasmonate (MeJA) (Chen et al., 2005). The overexpression of myo-inositol-1-phosphate synthase (*IbMIPS1*) in sweet potato has been shown to significantly upregulate the photosphoribulokinase (*PRK*) gene, thereby enhancing the plant’s salt tolerance (WANG et al., 2016). In soybeans, the expression of the stress-induced RD22-like protein (*GmRD22*) can mitigate salt and osmotic stress (Wang et al., 2012). Similarly, the grapevine *RD22* gene has been found to confer drought resistance when expressed in tobacco (Jardak-Jamoussi et al., 2016). Mitogen-activated protein kinase (MAPK) cascades play crucial roles in responding to both biotic and abiotic stresses, including drought. Notably, *SlMAPK1* is activated by various abiotic stresses and hormone treatments, and its overexpression in transgenic tomato plants has been linked to improved drought tolerance (Wang et al., 2018). *VvCCoAOMT* is a multifunctional O-methyltransferase, potentially playing an important role in the methylation of anthocyanins in grape berries, particularly under conditions of drought stress (Giordano et al., 2016). Cysteine synthase (CS) is an enzyme that catalyzes the biosynthesis of cysteine in plants. Cysteine serves as a precursor for the synthesis of various sulfur-containing metabolites, the most important of which is glutathione, which is used as a universal antioxidant and detoxifying agent to cope with various stresses (Yang et al., 2007). Overexpression of *CS* has been shown to significantly enhance the tolerance of tobacco plants to heavy metals and sulfur containing pollutants (Noji et al., 2001; Kawashima et al., 2004). In our study, we observed that drought stress induces the expression of *HcERF5* (Fig. 2), and that HcERF5 interact with various proteins, including HcPRK, HcRD22, HcMAP2, HcCS, HcCCoAOMT3, among others, as demonstrated by the yeast two-hybrid system (Fig. 8, Table S2). The expression of these proteins in *HcERF5*-silenced kenaf plants had significantly changed after silencing under drought stress treatment (Fig. 9), suggesting that *HcERF5* was involved in stress signal transduction and transcriptional regulation of downstream genes. In addition, the chlorophyll a-b binding protein is an important product of chlorophyll synthesis and photosynthesis. Overexpression of apple *MdLhcb4.3* in *Arabidopsis* was shown to significantly improve chlorophyll content and drought resistance of plants (Zhao et al., 2020). In Paeonia ostia, *PoWRKY71* directly binds to the W-box element of the *PoCAB151* promoter, thereby activating its transcription. Furthermore, plants overexpressing *PoCAB151* exhibited enhanced drought resistance, increased chlorophyll content, and improved photosynthetic activity, compare to those of *PoCAB151*-silenced plants (Luan et al., 2023). *OsERF19* plays an active role in salt stress and ABA response in rice and that *OsERF19* can directly target promoters of *OsOTS1* and *OsNCED5* and further increase their transcription level (Huang et al., 2021). We further confirmed the interaction between HcERF5 and HcCAB in the nucleus using BIFC technology (Fig. 10). Therefore, we hypothesized that *HcEFR5* may bind to specific cis-acting elements of the *HcCAB* promoter to regulate *HcCAB* expression and thus participate in response to drought stress. Further work will focus on the screening and identifying the cis-acting elements of *HcEFR5* that specifically bind to *HcCAB* promoters by experimental means such as Co-IP and EMSA, thereby regulating their expression and further revealing their molecular mechanisms in response to drought stress.

**Fig. 10.**
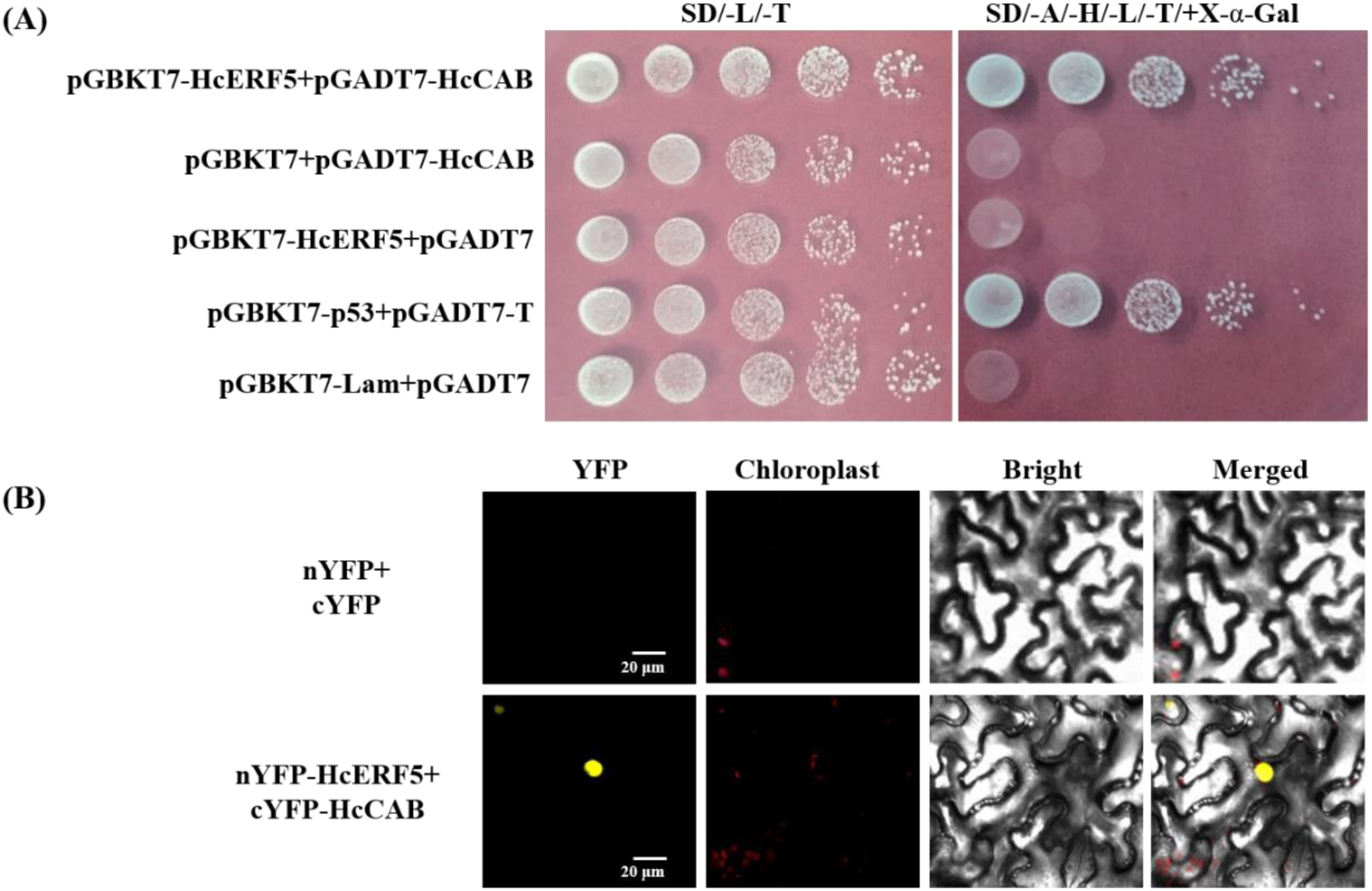
Interaction verification assay of HcERF5 and HcCAB proteins. (A) Validation of HcERF5 and HcCAB proteins using yeast two-hybrid assay. (B) Interaction between HcERF5 and HcCAB verified by BIFC system. Bar=20 μm

**Fig. 11.**
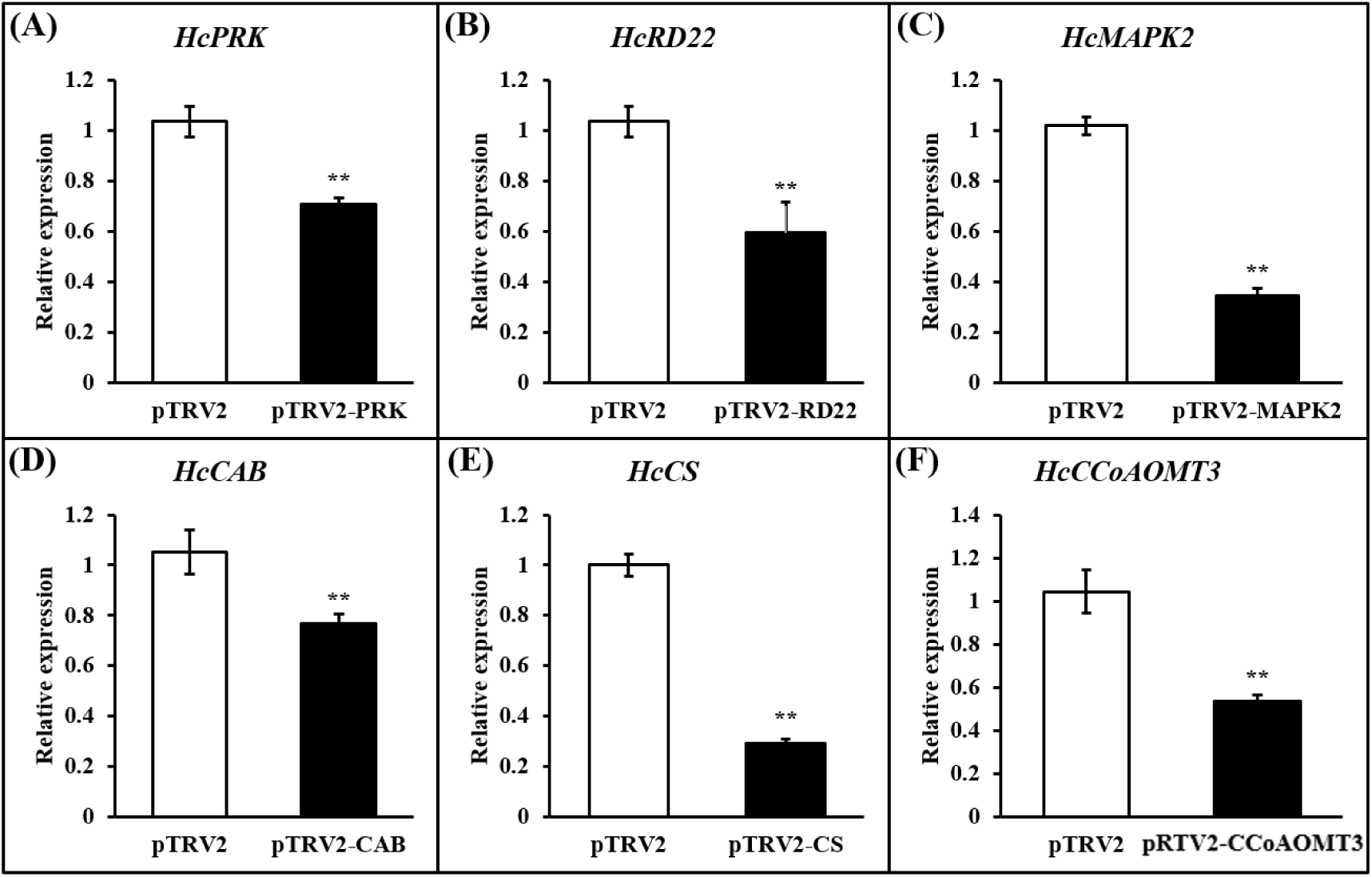
Expression profile of stress-responsive genes (A-H: *HcPRK*, *HcRD22*, *HcMAPK2*, *HcCAB*, *HcCS*, and *HcCCoAOMT3*) in *HcERF5*-silenced kenaf plants. Asterisks indicate statistical significance (* for *p* < 0.05 and ** for *p* < 0.01).

## CONCLUSION

This study systematically investigates the role and mechanism of *HcERF5* in regulating drought stress in kenaf. *HcERF5* expression is significantly induced by drought and ABA. Mutation of the *aterf5* gene in *Arabidopsis* significantly increased the sensitivity of seeds and seedlings to drought, whereas overexpression of *HcERF5* enhanced their drought stress tolerance. Silencing *HcERF5* in kenaf resulted in increased ROS levels and decreased ABA content, leading to poor drought tolerance, thus indicating the critical role of *HcERF5* in kenaf’s drought resistance. Moreover, six stress-responsive genes were identified to interact with HcERF5 via yeast Y2H assays, all of which were down-regulated in *HcERF5*-silenced kenaf plants under drought conditions compared to WT. RNA-seq revealed 2,489 DEGs in *HcERF5*-silenced plants, including genes involved in the ABA signaling pathway and antioxidant enzyme activity. A proposed regulatory model suggests that *HcERF5* enhances drought tolerance in kenaf by improving ROS scavenging abilities through the regulation of ABA synthesis and key stress resistance genes (Fig. 12). This study is of great significance for understanding the mechanism of *HcERF5* regulation in kenaf drought stress.

**Fig. 12.**
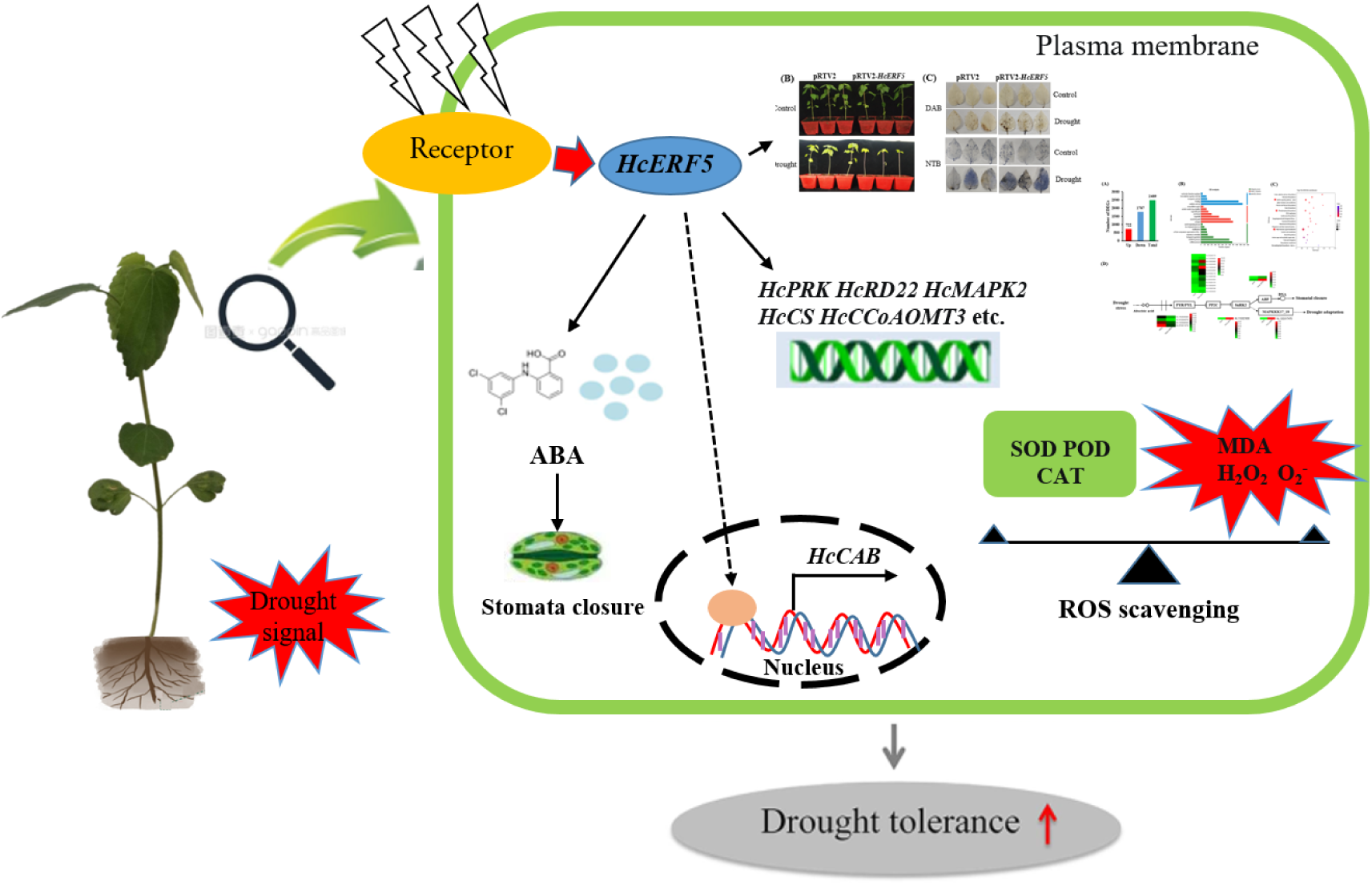
A proposed model of *HcERF5* regulating drought tolerance in kenaf.

## MATERIALS AND METHODS

### Identification, sequence analysis and subcellular localization of HcERF5

The amino acid sequence of the HcERF5 gene was predicted using DNAMAN 8.0 and Jalview software. The NCBI BLAST tool (https://www.ncbi.nlm.nih.gov/Structure/cdd/wrpsb.cgi) was used to perform conserved domain analysis and homology comparison study. The Phyre2 server (http://www.sbg.bio.ic.ac.uk/∼phyre2/html/page.cgi?id=index) was utilized to predict the three-dimensional structures and transmembrane regions. The neighbour-joining (NJ) method was employed to construct the phylogenetic tree in MEGA 6.06. The pBI121-HcERF5-EGFP vector, under the control of the CaMV 35S promoter (Fig. S1), was employed to examine the subcellular localization of the HcERF5-GFP fusion construct, following the detailed protocol outlined by Yue et al. (2022). The primers used in this study areprovided in Table S1.

### Expression analysis of the *HcERF5* and GUS staining

Total RNA was extracted from kenaf leaves treated with 20% PEG6000 and 100 μM ABA for various time points (0, 2, 6, 12, 24, and 48 hours) using TRIzol reagent (Vazyme Biotech Co., Ltd), following the manufacturer’s protocol. Additionally, total RNA was isolated from the leaves, petioles, stems, and roots of naturally growing kenaf seedlings. First-strand cDNA synthesis was carried out using a reverse transcription kit from Vazyme Biotech Co., Ltd, and total RNA. Quantitative real-time PCR (qRT-PCR) was caried out in a Bio-Rad CFX-96 RT-PCR amplifier (Bio-Rad, USA) using Cham SYBR qPCR master mix (Vazyme Biotech Co., Ltd, Nanjing, China). The 2^-ΔΔCT^ method was employed to analyze the relative expression levels, with the Actin3 gene serving as an internal reference. Table S1 contains a list of the qRT-PCR primers utilized in this investigation.

A 1.5 kilobase region upstream of the start codon ATG of the *HcERF5* gene was extracted and utilized to control the expression of the *HcERF5* gene in the binary expression vector pCambia1391Z. The constructed vector was transferred into *Agrobacterium tumefaciens* GV3101 using the freeze-thaw method, and then transgenic strain was introduced into Arabidopsis thaliana via the floral-dip technique. Following stress treatment, the positive seedlings were subjected by GUS staining. Subsequently, the seedlings were decolorized in a 75% (v/v) ethanol solution to remove chlorophyll before being photographed.

### Abiotic stress assays

In this study, wild-type (WT), *aterf5* mutants, and *HcERF5*-overexpressing (*HcERF5*-OE) *Arabidopsis* lines were used for abiotic stress analysis. For overexpressing *HcERF5*-OE lines, *Agrobacterium tumefaciens* GV3101 strain containing the full-length coding sequence of *HcERF5* was introduced into WT *Arabidopsis thaliana* through the floral-dip method and were grown until T3 generation. T-DNA inserted *aterf5* mutant (SALK_208574) seed were obtained from the *Arabidopsis* Biological Resources Centre. For stress analysis, the surface sterilized seeds of WT, *aterf5* mutants and *HcERF5*-OE were sown on the half-strength MS medium or a mixture of peat moss, vermiculite, and perlite in a 3:1:1 ratio. The plants were grown at 23 ± 2°C under long day conditions, with 16 hours of light and 8 hours of darkness. After 1 week, *Arabidopsis* seedlings were initially grown vertically on half-strength MS plates and then transferred to new plates containing mannitol (200 or 400 mM) and ABA (2 or 4 µM) for root growth assay. The primary root lengths were measured seven days after transplantation. For drought stress analysis on *HcERF5*-OE *Arabidopsis* plants, seven-day-old seedlings grown on half-strength MS plates were transferred to 7 cm square pots containing a mix of peat moss, organic substrate, and vermiculite in a 2:2:1 ratio. Drought stress was induced by withholding water for seven days, followed by a three-day rehydration period. Measurements of chlorophyll content, fresh weight, survival rate, and relative water content (RWC) were taken for WT, *aterf5* mutants, and *HcERF*5-OE seedlings. RWC was calculated using the formula: RWC (%) = [(FW − DW)/(TW − DW)] × 100, where fresh weight (FW), turgid weight (TW, after a 6-hour incubation in distilled water at room temperature), and dry weight (DW) were recorded.

### VIGS

The kenaf cultivar ‘Fuhong 992’ (FH992) was used to explore the role of the HcERF5 gene in response to drought stress using virus-induced gene silencing (VIGS). The SGN-VIGS web tool (https://vigs.solgenomics.net/) was used to identify the optimal target and design primers. The primer sequences can be obtained in Table S1, whereas the precise positions for amplification are marked in Fig. S6. VIGS derived tobacco rattle virus (TRV) vectors, pTRV1 and pTRV2 were used for further analysis. To create pTRV2-*HcERF5*, a partial fragment of the *HcERF5* gene was cloned into the pTRV2 vector. Then the auxiliary plasmid pTRV1, the empty plasmid pTRV2, and the positive recombinant plasmid pTRV2-*HcERF5* were successfully transformed into the *Agrobacterium* strain GV3101. At the first true leaf stage, the leaves of FH992 were injected with a cell suspension of the *Agrobacterium* (Luo et al., 2023). After being exposed to darkness for twenty-four hours, kenaf seedlings were then grown normally for 10 days. To assess the efficacy of gene silencing, random samples of kenaf plants were taken at the third true leaf stage and subjected to qRT-PCR analysis. The VIGS seedlings that tested positive were then exposed to a 10-day natural drought stress in the soil and the growth and physiological indicators were examined accordingly.

The leaves of kenaf plants were examined with an inverted NICON microscope with a 40X objective lens, and structural analysis was conducted with ProgRes. The guard cell located on the lower epidermis of the leaves was captured using ProgRes® CapturePro 2.8.8 software. The stomatal aperture was measured using ImageJ software. The experiment was replicated thrice, with 50-60 stomata measured in each treatment group. The density of stomata was obtained by calculating the number of pores in each photo/the area of the photo (204.8×153.6 μm^2^), with 10 photos used for statistical analysis

The content of hydrogen peroxide (H_2_O_2_), superoxide anion radical (O_2_^-^) and proline content were measured according to the method described by Luo et al. (2023) (Luo et al., 2023). Malondialdehyde (MDA) content and the activities of superoxide dismutase (SOD), peroxidase (POD), catalase (CAT) and glutathione reductase (GR) were measured according to the method described in our previous study (Chen et al., 2020). The concentrations of hydrogen peroxide (H_2_O_2_) and superoxide radicals (O_2_^-^) were determined by staining with 3, 30-diaminobenzidine (DAB) and nitroblue tetrazolium (NBT), respectively. The ABA content was determined by LC-MS (Agilent 1290 infinity-SCIEX B5000trap, https://www.agilent.com/).

For the water loss rate assay, the true leaves of pTRV2 and pTRV2-*HcERF*5 plants were used as material. The intact leaves were cut off to measure fresh weight (FW) immediately. When measuring the water loss rate of leaves, the isolated leaves were placed in a greenhouse to ensure that the environment of the isolated leaves was the same as that of the experimental and control plants at intervals of 0.5 h, 1 h, 1.5 h, 2 h, 3 h and 5 h. The weight of the isolated blade W0, and calculate the final result according to the formula of water loss rate (%) = (W0-FW)/W0.

### Transcriptome sequencing

The leaves of kenaf seedlings in pTRV2 and pTRV2-HcERF5 plants under drought stress were collected for transcriptome sequencing, and each treatment contained three biological replicates. RNA extraction, library construction, sequencing, and data processing were conducted by Majorbio (Shanghai, China) based on Illumina Novaseq 6,000 sequencing platform. The clean data was then mapped to the reference genome sequence (https://bigd.big.ac.cn/gwh/Assembly/1033/show). The expression levels of genes/transcripts were quantitatively analyzed usimg Fragments Per Kilobases per Millionreads (FPKM). DEseq2 software was employed to search for differentially expressed genes (DEGs) between two different treatment groups. The p-value < 0.05 and |log_2_ fold change| ≥1 was considered as the threshold of DEGs. Functional analysis of DEGs was performed via Gene Ontology (GO), and Kyoto Encyclopedia of Genes and Genomes (KEGG) on the free online platform of Majorbio (www.majorbio.com).

### Yeast two-hybrid (Y2H) and bimolecular fluorescence complementation (BIFC) assay

Specific cloning primers were designed based on the sequence of the CDS of the *HcERF5* gene (Table S1), and seamlessly cloned into the bait vector pGBKT7. The recombinant plasmids pGBKT7-HcERF5, empty PGBKT7 plasmids (negative control), and pGBKT7-53+pGADT7-T (positive control) were transformed into the Y2H Gold strain using the LiTE/PEG method. The monoclonal yeast was identified by PCR and then the bacterial solution was diluted 1, 10, 100, and 1000 folds. the diluted sample (2 μL) was spread on SD/-Trp board and SD/-Trp-Leu-His+X-α-gal (SD/-TDO+X-α-gal) plate and cultured at 30 ℃ for 3-5 days for validation of toxicity and self-activation.

For initial screening of HcERF5 protein, the bait vector pGBKT7-HcERF5 with library mixture (matching) was used on SD/-Trp-Leu-His medium, and selected monoclonal clones were spread on SD/-Trp-Leu+X-α-Gal (SD/-DDO+X-α-Gal) medium. The blue spot colony was amplified and sequenced with T7 and ADF/B primers (Table S1) and the sequencing results were aligned against the kenaf genome (https://bigd.big.ac.cn/gwh) (Zhang et al., 2020) and NCBI BLAST (https://blast.ncbi.nlm.nih.gov). Finally, the bacterial cultures of positive clones, pGBKT7-53+pGADT7-T (positive control), and pGBKT7-HcERF5+pGADT7 (negative control) were cultured to OD=0.5, and spread 3 μL onto SD/-Trp-Leu and SD/-Trp-Leu-His-Ade+X-α-Gal (SD/-QDO+X-α-Gal) medium plates. The plates were then incubateed at 30 ℃ for 3 days and photographed.

The coding sequences of *HcERF5* and *HcCAB* were cloned into the vectors pCAMBIA1301-nYFP and pCAMBIA1301-cYFP, and transferred into *Agrobacterium* GV3101, which can be immediately expressed in tobacco. The fluorescence of YFP was analyzed by confocal laser scanning microscope (TCS-SP8MP; Leica, Germany). The flesh cell on tobacco leaf were observed by adjusting the excitation wavelength at 488 nm and emission wavelength at 507 nm.

### Statistical analysis

Excel 2019 was used to organize the data and GraphPad Prism 8 software was used for plotting. SPSS v24.0 software was used for analysis of variance, and Duncan’s new complex range method was used to compare the significance of differences between treatments (* for *p* < 0.05 and ** for *p* < 0.01).

## Supplemental Data

**Supplemental Table S1 Primers for vector construction, qRT-PCR, and VIGS**

**Supplemental Table S2 The proteins of interaction with HcERF5**

**Supplemental Table S3 List of DEGs**

**Supplemental Table S4 GO analysis of DEGs**

**Supplemental Table S5 KEGG analysis of DEGs**

## Abbreviations

GUS: β -glucuronidase
ABA: abscisic acid
PEG: polyethylene glycol
MS: Murashige and Skoog
OE: overexpressing
GFP: green fluorescent protein
ROS: reactive oxygen species
VIGS: Virus-induced gene silencing
WT: wild type
CAT: Catalase
MDA: malondialdehyde
POD: peroxidase
SOD: superoxide dismutase
qRT-PCR: quantitative real-time PCR
DAB: 3, 3′ -diaminobenzidine
NBT: nitroblue tetrazolium
H_2_O_2_: hydrogen peroxide
O_2_^-^: superoxide anion radical

